# Extent of the annual Gulf of Mexico hypoxic zone influences microbial community structure

**DOI:** 10.1101/483735

**Authors:** Lauren Gillies Campbell, J. Cameron Thrash, Nancy N. Rabalais, Olivia U. Mason

## Abstract

Rich geochemical datasets generated over the past 30 years have provided fine-scale resolution on the northern Gulf of Mexico (nGOM) coastal hypoxic (≤ 2 mg of O_2_ L^-1^) zone. In contrast, little is known about microbial community structure and activity in the hypoxic zone despite the implication that microbial respiration is responsible for forming low dissolved oxygen (DO) conditioXSns. Here, we hypothesized that the extent of the hypoxic zone is a driver in determining microbial community structure, and in particular, the abundance of ammonia-oxidizing archaea (AOA). Samples collected across the shelf for two consecutive hypoxic seasons in July 2013 and 2014 were analyzed using 16S rRNA gene sequencing, oligotyping, microbial co-occurrence analysis and quantification of thaumarchaeal 16S rRNA and archaeal ammonia-monooxygenase (*amoA*) genes. In 2014 Thaumarchaeota were enriched and inversely correlated with DO while Cyanobacteria, Acidimicrobiia and Proteobacteria where more abundant in oxic samples compared to hypoxic. Oligotyping analysis of *Nitrosopumilus* 16S rRNA gene sequences revealed that one oligotype was significantly inversely correlated with dissolved oxygen (DO) in both years and that low DO concentrations, and the high Thaumarchaeota abundances, influenced microbial co-occurrence patterns. Taken together, the data demonstrated that the extent of hypoxic conditions could potentially influence patterns in microbial community structure, with two years of data revealing that the annual nGOM hypoxic zone is emerging as a low DO adapted AOA hotspot.

## Introduction

Deoxygenation of the ocean is one of the primary consequences of global climate change [1,2], with much attention directed towards coastal hypoxic zones (dissolved oxygen (DO) concentrations below 2 mg L^-1^ or 62.5 μmol/kg). Hypoxic zones are frequently referred to as “dead zones” because they are inhospitable to macrofauna and megafauna; however, microorganisms thrive in such environments [3]. Eutrophication-associated dead zones have been reported in over 500 locations spanning the globe [4] and are predicted to increase in number and size in the near future as a result of increasing greenhouse gas emissions [2]. The northern Gulf of Mexico (nGOM) is the site of the second largest eutrophication-associated coastal dead zone in the world, with bottom water hypoxia extending to over 20,000 km^2^ [5] and covering anywhere from 20% to 50% of the water column during the summer months [6]. The nGOM hypoxic zone is influenced by the freshwater input and nutrient load from the Mississippi (MI) and Atchafalaya (AR) Rivers [7], which results in a phytoplankton bloom, the biomass of which is subsequently respired by aerobic microorganisms leading to low DO concentrations or hypoxic zones [8,9]. Hypoxia in the nGOM has increased in severity during the summer months in direct response to additional inorganic nitrogen loading in the Mississippi watershed, beginning around the 1950s, after which the nitrate flux to the nGOM continental shelf tripled [10-12]. Therefore, thermal warming, nutrient rich freshwater discharge and microbial respiration culminate and result in an annually extensive nGOM hypoxic zone, that reached a record size at 8,776 mi^2^ (22,720 km^2^) in 2017 (LUMCON 2017, https://gulfhypoxia.net/research/shelfwide-cruise/?y=2017, accessed 1/22/2018).

While the sequence of events that leads to the nGOM hypoxic zone are well documented [4,13-16], efforts to understand how these chemical, biological, and physical factors influence microbial community structure, abundances, and activity across environmental gradients in the nGOM coastal shelf have only recently been undertaken [17-21]. King and colleagues [17] reported that before the onset of hypoxia (March), Alpha- and Gamma- Proteobacteria, Bacteriodetes and Actinobacteria are abundant in nGOM waters <100m, with Planctomycetes and Verrucomicrobia being less abundant. Gillies and colleagues [20] sampled during the 2013 hypoxic event in late July, when hypoxic conditions historically and predictably prevail, and reported that the high normalized abundance of a Thaumarchaeota (100% similar to the ammonia-oxidizing archaea *Nitrosopumilus maritimus* [22]) Operational Taxonomic Unit (OTU) (16S rRNA gene iTag data) and the absolute abundance of Thaumarchaeota 16S rRNA and archaeal *amoA* gene copies (quantitative PCR (qPCR)) were significantly inversely correlated with DO [20]. In this 2013 study, Thaumarchaeota comprised more than 40% of the microbial community (iTag sequence data) in hypoxic samples [20]. In contrast, King and colleagues [17] and Tolar and colleagues [23] showed Thaumarchaeota were present at depths <100m increasing in abundance at depths >100m in the nGOM when conditions are oxic over the shelf. Bristow and colleagues [19] sampled a similar geographic area in 2012 during a historically small hypoxic zone (smallest recorded since 1988) and reported that thaumarchaeal abundances increased with depth; reaching a maximum at 120 m. In their shallow, bottom water (15 m) sample, Thaumarchaeota reached ~15% of the microbial community and the highest rate of ammonia oxidation were reported at this single hypoxic station ([19]; Station 6, see their Fig 3A). Further, genomic reconstruction of two *Nitrosopumilus* genomes (79% and 96% complete) assembled from shotgun metagenomic data from a subset of the 2013 hypoxic zone samples [20], combined with complimentary shotgun metatranscriptomic data, revealed that the *Nitrosopumilus* reported in that study were active in the 2013 hypoxic zone (Campbell and Mason, unpublished). Specifically, transcripts from *amoABC* genes, with the same synteny as observed in the genome of *N. maritimus* [24] were identified. These data suggest that nitrification, particularly ammonia oxidation, was actively being carried out in the 2013 hypoxic zone by Thaumarchaeota-- an aerobic process that would continue to draw down oxygen, exacerbating hypoxic conditions across the shallow continental shelf [25,26].

**Figure 3:**
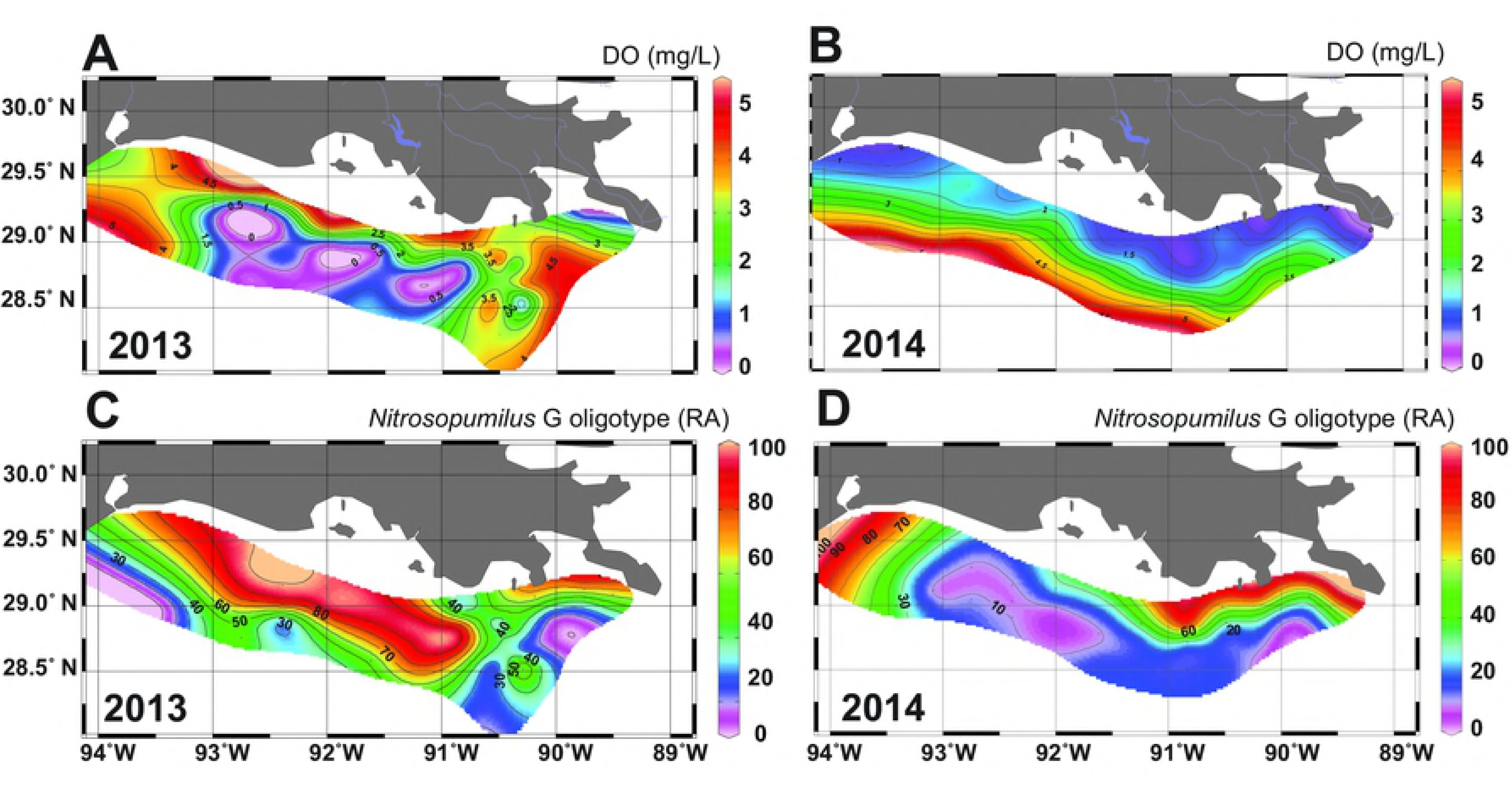
Ocean Data View plots of DO and *Nitrosopumilus* G oligotype data. Plots of **(A)** DO concentrations for the same samples collected in Y13 **(B)** and Y14. **(C)** Oligotype data for *Nitrosopumilus* G oligotype for the same samples collected in Y13 (**C**) and Y14 **(D)**.

In the nGOM hypoxic zone, the importance of AOA is only beginning to emerge; however, AOA have been shown to be more abundant than ammonia-oxidizing bacteria (AOB) in terrestrial and marine ecosystems, [27-31], and it is proposed that AOA can outcompete bacteria in low energy conditions [32]. In other oxygen minimum zones (OMZs), several studies have reported an increase in abundance of Thaumarchaeota in low DO samples [18,33-37]. The abundance of archaeal *amoA* transcripts also increases in OMZs [35,36,38].

Therefore, to begin to fill in the knowledge gaps on microbial community structure and specifically on AOA abundances in the nGOM hypoxic zone, we describe the microbial communities in samples collected in July 2014 within and outside of the 13,080 km^2^ hypoxic zone, and compared these to Y14 results to the same thirty-three sites sampled during the Y13 hypoxic zone (15,120 km^2^) [20]. We expanded sampling in Y14 to include surface water and more bottom water oxygen minimum zone samples, as compared to Y13. Our goal was to determine the influence the extent of the hypoxic zone has on the overall microbial ecology and in particular the abundance of AOA across hypoxic seasons in the nGOM. Further, we sought to determine if the high AOA abundances observed in low DO conditions in this large coastal hypoxic zone occur predictably and, if so, consider the potential ecological implications for an annual increase of AOA in the hypoxic zone.

## Material and Methods

### Sample collection

The 2014 hypoxic zone was mapped over seven days, from 27 July to 2 August 2014 and measured 13,080 km^2^. At each of the 52 sites sampled, a sample was collected at the surface (except for sites A’2, B9 and C7) (1 meter below sea level (mbsl); samples designated S for surface; 47 samples total) and near the seafloor at the oxygen minimum zone (19 mbsl avg. collection depth; samples designated B for bottom; 50 samples total). Samples were collected from the surface of the Mississippi River at two sites (0mbsl; designated R2 and R4 for Mississippi River) for a total of ninety-nine samples. For each of the ninety-nine samples collected, temperature, depth, salinity (conductivity) and *in situ* chemistry were determined using a conductivity-temperature-depth (CTD) instrument (SeaBird SBE32 5L bottle carousel). Concentrations of DO, ammonium (NH_4_), nitrite (NO_2_) + nitrate (NO_3_), phosphate (PO_4_) and chlorophyll a were determined. The sample location map and subsequent plot of DO data were made with Ocean Data View [39].

### Oxygen, chlorophyll a and nutrients

Oxygen concentrations were determined *in situ* with the CTD dissolved oxygen sensor. Oxygen concentrations were verified using Winkler titrations [40] shipboard. Chlorophyll a samples were concentrated on 25-mm Whatman GF/F filters from 500 ml ^−1^ L seawater and stored at −20°C.

Chlorophyll a was extracted using the methods described in the Environmental Protection Agency Method 445.0, “In Vivo Determination of Chlorophyll a in Marine and Freshwater Algae by Fluorescence” [41]; however, no mechanical tissue grinder or HCl were used.

Chlorophyll a concentrations were determined using a fluorometer with a chlorophyll a standard (*Anacystis nidulans* chlorophyll a). For nutrients, 60 ml of seawater was filtered through 0.22- μm Sterivex filters into two 30-ml Nalgene bottles and stored at −20°C. Nutrient concentrations were determined by the marine chemistry lab at the University of Washington following the WOCE Hydrographic Program using a Technicon AAII system (http://www.ocean.washington.edu/story/Marine+Chemistry+Laboratory).

### Microbial sampling and DNA extractions

From each station, up to 5 L of seawater were collected and filtered with a peristaltic pump both at the surface and at the oxygen minimum. A 2.7-μM Whatman GF/D pre-filter was used and samples were concentrated on 0.22-μM Sterivex filters (EMD Millipore). Sterivex filters were sparged, filled with RNAlater and frozen. Samples were transported to Florida State University on dry ice and stored at −80 until DNA extractions and purifications were carried out. DNA was extracted directly off of the filter by placing half of the Sterivex filter in a Lysing matrix E (LME) glass/zirconia/silica beads Tube (MP Biomedicals, Santa Ana, CA, USA) using the protocol described in [20] which combines phenol:chloroform:isoamyalcohol (25:24:1) and bead beating. Genomic DNA was purified using a QIAGEN (Valencia, CA, USA) AllPrep DNA/RNA Kit and quantified using a Qubit2.0 Fluorometer (Life Technologies, Grand Island, NY, USA).

### 16S rRNA gene sequencing and analysis

16S rRNA genes were amplified from 10 ng of purified genomic DNA in duplicate using archaeal and bacterial primers 515F and 806R, which target the V4 region of *Escherichia coli* in accordance with the protocol described in [42,43], used by the Earth Microbiome Project (http://www.earthmicrobiome.org/emp-standard-protocols/16s/), with a slight modification: the annealing temperature was modified to 60 °C. Duplicate PCR reactions were combined and purified using Agencourt AMPure XP PCR Purification beads (Beckman Coulter, Indianapolis, IN) and sequenced using the Illumina MiSeq platform. Ninety-nine samples from 52 stations were sequenced and analyzed. Raw sequences were joined using fastq-join [44] with the *join_paired_ends.py* command and then demultiplexed using *split_libraries_fastq.py* with the default parameters in QIIME version 1.9.1 [45]. Demultiplexed data matching Phi-X reads were removed using the SMALT 0.7.6 akutils phix_filtering with the *smalt map* command [46].

Chimeras were removed using *vsearch –uchime_denovo* with VSEARCH 1.1.1 [47]. Neither PhiX contamination nor chimeric sequences were observed. Sequences were then clustered into operational taxonomic units (OTUs), which was defined as ≥ 97% 16S rRNA gene sequence similarity using SUMACLUST [48]and SortMeRNA [49] with *pick_open_reference.py -m sortmerna_sumaclust*. Greengenes version 13.5 [50] was used for taxonomy. The resulting OTU table was filtered to keep only OTUs that had 10 sequences or more (resulting in 8,959 OTUs), and normalized using cumulative sum scaling (CSS) with metagenomeSeq [51] in R. These sequences will be available in NCBI’s SRA and on the Mason server at http://mason.eoas.fsu.edu. While taxonomy was determined using Greengenes, taxonomy for OTUs that were determined to be significantly (Wilcoxon) different between environments (surface and bottom, hypoxic and oxic, Y13 and Y14 samples) were additionally verified beyond the class level using SILVA ACT, version 132 alignment and classification [52] with the confidence threshold set to 70%. To determine close relatives of specific OTUs of interest NCBI’s nucleotide blastn was used to search the Nucleotide collection using Megablast with default parameters [53]. Pairwise sequence comparisons were also carried out using blastn.

### Statistics

Multiple rarefactions and subsequently the alpha diversity metrics Shannon (H’) [54], observed OTUs (calculates the number of distinct OTUs, or richness) and equitability (Shannon diversity/natural log of species richness; the scale is 0-1.0; with 1.0 indicating that all species are equally abundant) were calculated in QIIME version 1.9.1 using *multiple_rarefactions.py* followed by *alpha_diversity.py*. The Shapiro-Wilk test (*shapiro.test* function) was used to test diversity values and environmental variables for normality in R. In R, the Wilcoxon Rank-Sum test (*wilcox.test* function) with the application of Benjamini-Hochberg’s (B-H) False Discovery Rate (FDR) (alpha = 0.05) [55] was used to test for significant differences in diversity values, environmental variables, absolute abundance data (Thaumarchaeota 16S rRNA and *amoA* gene copy numbers per L of seawater) and CSS normalized oligotype data between surface and bottom samples, oxic and hypoxic samples and Y13 and Y14.

Statistically significant differences in all CSS normalized OTU abundances between surface and bottom samples, hypoxic and oxic conditions and between Y13 and Y14 samples were determined using the non-parametric Wilcoxon test in METAGENassist [56], with the B-H correction for multiple tests. Prior to the Wilcoxon test, data filtering was carried out to remove OTUs that had zero abundance in 50% of samples and the interquantile range estimate was used to filter by variance to detect near constant variables throughout [57]. After quality filtering 528 OTUs remained out of 8,960 OTUs for Y14 surface and bottom samples, 623 OTUs remained out of 7,924 OTUs for Y14 bottom only samples, and 724 OTUs remained out of 9,784 OTUs for the same samples collected in Y13 and Y14.

Beta-diversity of CSS normalized data was examined using non-metric multidimensional (NMDS) scaling with Bray-Curtis in R with the *metaMDS* function in the vegan package [58]. The *envfit* function in vegan was then used to fit vectors of environmental parameters onto the ordinations with p-values derived from 999 permutations with the application of the B-H correction for multiple tests using the *p.adjust* function (we defined corrected p-values ≤ 0.05 as significant). Using the vegan package in RStudio, the non-parametric test adonis [59] was used to test whether microbial community composition was significantly different between clusters of samples (surface and bottom, hypoxic and oxic, year 2013 and 2014, east and west latitudes). The *betadisper* function (vegan) was used to test for homogeneity of multivariate dispersion among sample clusters for surface/bottom samples, hypoxic/oxic samples, Y13/Y14 samples and east/west samples. Betadisper p-values were derived from 999 permutations using the *permutest* function and a B-H correction for multiple tests was performed. Using the psych package [60] in R, the nonparametric Spearman’s rank order correlation coefficient (rho (ρ)) and p-values (B-H correction) were determined for environmental variables and CSS normalized OTUs that were significantly different (Wilcoxon) for Y14 surface and bottom samples, hypoxic and oxic samples and Y14 and Y13 same samples.

Co-occurrence analysis between OTUs that were significantly (Wilcoxon) different between Y13 and Y14 was carried out by determining Spearman’s correlation coefficients using the psych package in R, similar to [61-64]. Spearman’s correlation results were visualized in R for OTUs that were significantly (corrected p-values ≤ 0.05) correlated with one or more of the three Thaumarchaeota OTUs (4369009, 1584736 and 4073697).

### Oligotyping

Shannon entropy (oligotyping) [65,66] analysis was carried out on all 16S rRNA gene sequences identified as *Nitrosopumilus* to identify variability in specific nucleotide positions in this genus. Scripts were used to format QIIME generated data for oligotyping using *q2oligo.py* and *stripMeta.py* [67]. The QIIME command *filter_fasta.py* was used to obtain all *Nitrosopumilus* 16S rRNA gene sequences. All *Nitrosopumilus* 16S rRNA gene data was then analyzed using the oligotyping pipeline version 0.6 for Illumina data [65,66] with the following commands, *o-trim*, *o-pad-with-gaps* and *entropy-analysis*. All oligotype data was CSS normalized using metagenomeSeq in R.

### Quantitative PCR

Thaumarchaeal and bacterial 16S rRNA and archaeal *amoA* genes were quantified in duplicate using the quantitative polymerase chain reaction (qPCR) assay. For each qPCR reaction 10 ng of genomic DNA was used. Thaumarchaeota 16S rRNA genes were amplified using 334F and 554R with an annealing temperature of 59°C [68]. Bacterial 16S rRNA genes were amplified using 1369F and 1492R with 56°C as the annealing temperature [68]. Archaeal *amoA* genes were amplified using Arch-*amoA*-for and Arch-*amoA*-rev with 58.5°C as the annealing temperature [29]. Standards (DNA cloned from our samples for Thaumarchaeota 16S rRNA and archaeal *amoA* and *E. coli* for bacterial 16S rRNA) were linearized, purified, and quantified by fluorometry. The reaction efficiencies for the standard curve were calculated from the slope of the curve for all qPCR assays and were 91.1% for Thaumarchaeota 16S rRNA genes, 88.5% for bacterial 16S rRNA genes and 85.3% for archaeal *amoA* genes. The qPCR data was converted to gene copies L^-1^ of seawater.

## Data deposition

All sequences reported in this paper will be deposited into the NCBI sequence archive upon article acceptance to the journal.

## Results

### In situ chemistry and physical attributes of the 2014 hypoxic zone

All environmental parameters measured, including DO concentrations in all ninety-nine surface and bottom samples and the two surface samples from sites at the MI River mouth (R4 and R2) collected in late July from Y14 are shown in Supplementary Table 1. In Y14, all environmental variables (S1 Table) except chlorophyll a were significant between surface and bottom samples (Wilcoxon with B-H corrected p-values; S2 Table). Average NO_2_ + NO_3_ concentrations were higher in surface samples compared to bottom samples, while NH_4_ and PO_4_ were higher in bottom samples.

Bottom water hypoxic conditions in 2014 were confined to the coast and shallower depths (S1 Fig), reaching a total area of 13,080 km^2^. When looking at bottom samples only, all environmental variables except NH_4_, temperature and chlorophyll a were significantly different between hypoxic and oxic samples (S2 Table). Average NO_2_+NO_3_ and PO_4_ concentrations were higher in hypoxic water samples compared to oxic samples, while average salinity concentrations were higher in oxic samples. The average depth of the bottom water hypoxic samples was 15m and the average depth of oxic samples was 25m.

### Microbial community composition across the shelf and with depth in the 2014 hypoxic zone

ITag sequencing of 16S rRNA genes was used to determine microbial community composition across the shelf in the Y14 nGOM hypoxic zone, surface and bottom samples. This sequencing effort resulted in 11.9 million reads and 8,959 number of OTUs (the full OTU table is included as S3 Table). In the two MI River samples, Actinobacteria, Cyanobacteria and Proteobacteria were the most abundant phyla (Fig 1A) (Thaumarchaeota relative abundances were low, with R4 relative abundances being 1.4% and R2 abundances <0.001%). Actinobacteria OTU4345058 (ac1) relative abundance was the highest (up to 22% relative abundance at site R4 and 0.7% at the more saline R2 site), decreasing to 0.1% in surface samples near the mouth of the MI River to < 0.001% to undetectable outside of the MI River in surface and bottom samples (avg. in surface samples 3.54 × 10^-4^ and avg. in bottom samples 3.8 × 10^-5^). Cyanobacteria (*Cyanobium* PCC-6307) OTU404788 was the second most abundant OTU in river samples with a higher abundance in the more saline R2 sample (up to 28% at site R2 and 3% at R4). The surface microbial communities had similar dominant phyla to the MI River samples with Cyanobacteria (avg. 33%) and Proteobacteria (avg. 32%), predominantly Gamma- and Alphaproteobacteria being the most abundant, but had lower abundances of Actinobacteria (3%) than in the two river samples (Fig 1A). In surface samples the average relative abundance of Thaumarchaeota, *N. maritimus* OTU4369009 was 0.6%.

**Fig 1.**
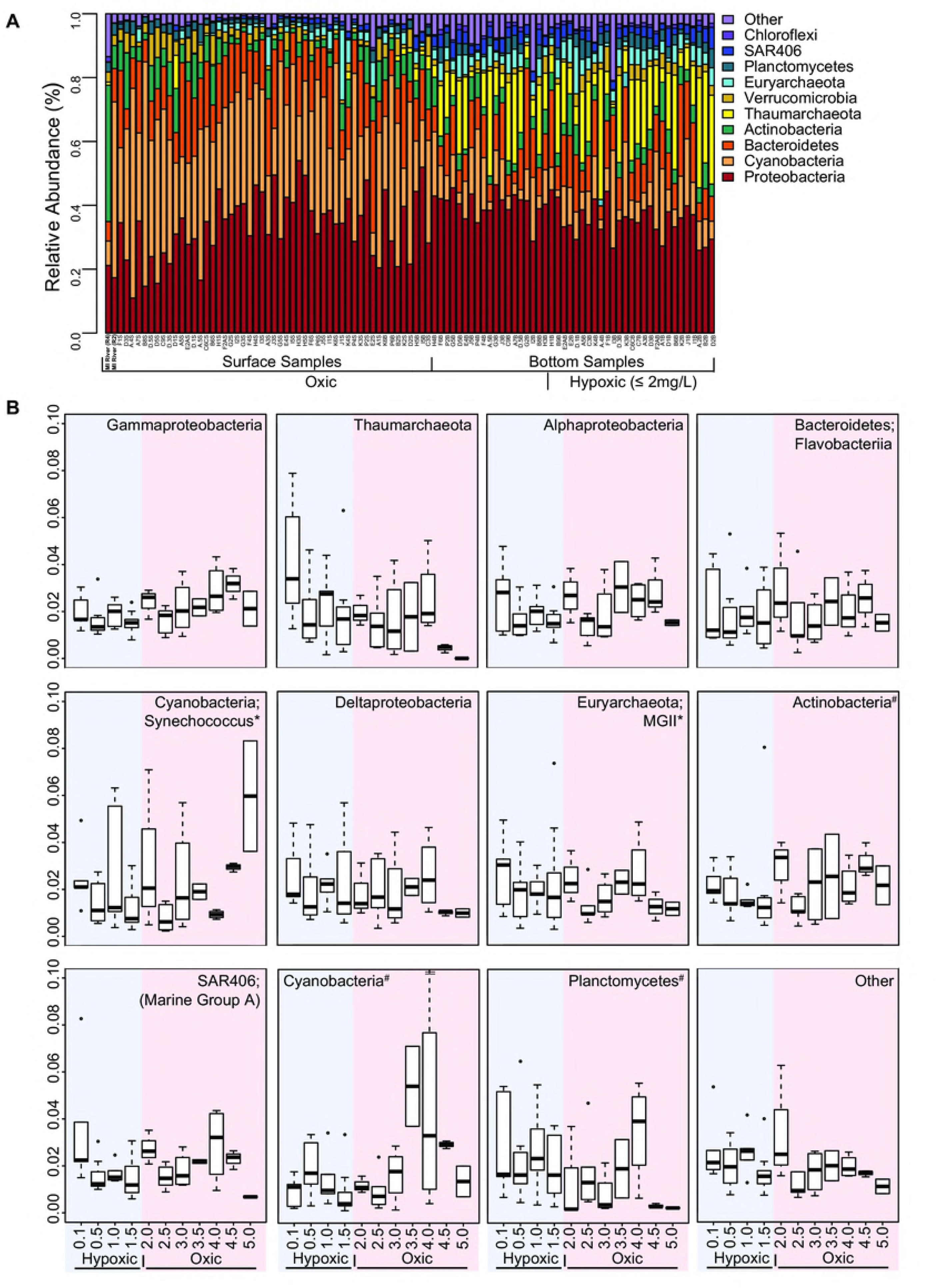
Phyla level bar graph and boxplots of most abundant bacterial and archaeal groups. **(A)** Phyla level bar graph of relativized 16S rRNA gene iTag sequence data, in which only the more abundant bacterial and archaeal groups are shown. Less abundant groups were summed under “Other.” Samples are sorted from lowest to highest DO concentrations on the x-axis and surface and bottom samples are differentiated by brackets with the two Mississippi (MI) River samples on the far left. **(B)** Boxplots of most abundant classes for bottom samples plotted along a DO gradient from lowest to highest DO concentrations (* indicates that taxonomy for these phyla were further refined based on the OTU taxonomy in these groups. # indicates that taxonomy was not resolved beyond phylum).

Bottom water (avg. collection depth was 19 m and avg. DO concentration was 2.26 mg L^-1^) samples were dominated by Proteobacteria (avg. 37%), primarily Gamma- Alpha- and Deltaproteobacteria, as well as Thaumarchaeota (avg. 14%) (Figs 1A and B). The dominant Thaumarchaeota, *N. maritimus* OTU4369009 had an average relative abundance of 13% in bottom water samples. When plotting bottom water data along a DO gradient, several trends emerged. Gammaproteobacteria, Alphaproteobacteria, Bacteroidetes, Cyanobacteria, and the Actinobacteria Acidimicrobiia generally increased in relative abundance with higher DO (Fig 1B). Deltaproteobacteria, MGII Euryarchaeota and Planctomycetes generally increased in abundance with decreasing DO (Fig 1B). In hypoxic samples, Thaumarchaeota were most abundant, particularly at the lowest DO concentration, with decreasing abundances as DO increased (Fig 1B). Of these taxa in the bottom samples, the normalized abundances of Thaumarchaeota and Planctomycetes were significantly inversely correlated with DO (Spearman’s ρ for Thaumaechaeota= −0.38 and Planctomycetes= −0.35, corrected p-values ≤ 0.05).

## Microbial ecology and correlation analyses with environmental variables across the 2014 hypoxic zone

### Surface and bottom samples

Statistical analysis of alpha diversity metrics revealed that microbial diversity (Shannon) was significantly lower in the surface samples when compared with bottom samples (Wilcoxon with B-H corrected p-values; S2 Table). Specifically, Shannon diversity indices averaged 5.97 in all nGOM surface samples and 6.02 in the two surface river samples, as compared with 6.45 in bottom samples. Richness (observed species) increased significantly with depth (avg. 329.92 in surface samples and avg. 494.89 in bottom samples). A test of evenness (equitability) between the surface and bottom sample types revealed that surface samples (avg. 0.72) were less even than bottom samples (avg. 0.75).

Non-parametric statistical analysis (Wilcoxon) was then used to determine which OTUs were responsible for the significant difference in species richness when comparing all surface and bottom samples. Seventeen OTUs showed significant differences in their CSS normalized abundances between surface and bottom samples (S2 Fig). Seven of these OTUs had higher average CSS normalized abundances in bottom samples and were significantly inversely correlated with DO (S2 Fig; Spearman’s ρ and corrected p-values ≤ 0.05 in S4 and 5 Tables). The dominant Thaumarchaeota, *N. maritimus* OTU4369009 was significantly inversely correlated with DO and positively correlated with NO_2_ + NO_3_ and PO_4_ (S4 and 5 Tables).

### Bottom water samples

When analyzing bottom water samples alone, Shannon diversity, richness and evenness indices for hypoxic versus oxic samples were not significantly different. However, comparison of the bottom hypoxic and oxic samples, including environmental variables (Wilcoxon), revealed that the CSS normalized abundances of 16 OTUs were significantly different depending on DO status (S3 Fig). Of the 16 OTUs, six had higher CSS normalized abundances in hypoxic samples and were significantly inversely correlated with DO (S3 Fig; Spearman’s ρ and corrected p-values ≤ 0.05 in S6 and 7 Tables). Thaumarchaeota, *N. maritimus* OTU4369009 comprised an average of 16% of the microbial community in hypoxic samples, with a peak abundance of 33% of the microbial community in sample A’2 bottom, which had one of the lowest DO concentrations (0.31 mg of O_2_ L^-1^), versus an average of 10% relative abundance in oxic samples. Further, Thaumarchaeota, *N. maritimus* OTU4369009 was significantly inversely correlated with DO in Y14 bottom samples and significantly inversely correlated with NH_4_ and temperature in bottom samples (S6 and 7 Tables). Thaumarchaeota, *N. maritimus* OTU4369009 was significantly positively correlated with NO_2_+NO_3_, PO_4_ and salinity in bottom samples (S6 and 7 Tables). While this Thaumarchaeota OTU4369009 was abundant in hypoxic samples and inversely correlated with DO, the normalized abundances were not significantly different between hypoxic and oxic bottom samples (Wilcoxon).

### Microbial community organizational structure and drivers in the 2014 hypoxic zone

To examine the primary drivers in structuring the 2014 microbial communities in the surface and in the bottom water nGOM hypoxic zone, whole community 16S rRNA gene sequence data were examined using Bray-Curtis distances with non-metric multidimensional scaling (NMDS). All environmental variables shown as vectors were significantly correlated (corrected p-values ≤ 0.05) with NMDS axes revealing the primary drivers in influencing the microbial community structure to be DO, depth, NO_2_+NO_3_, and PO_4_ (Fig 2 and S8 Table), with NO_2_+NO_3_ and PO_4_, decreasing with increasing distance from the MI river mouth. While the adonis test for all Y14 samples was significant, so too was beta-dispersion, suggesting non homogenous dispersion for these sample clusters.

**Figure 2.**
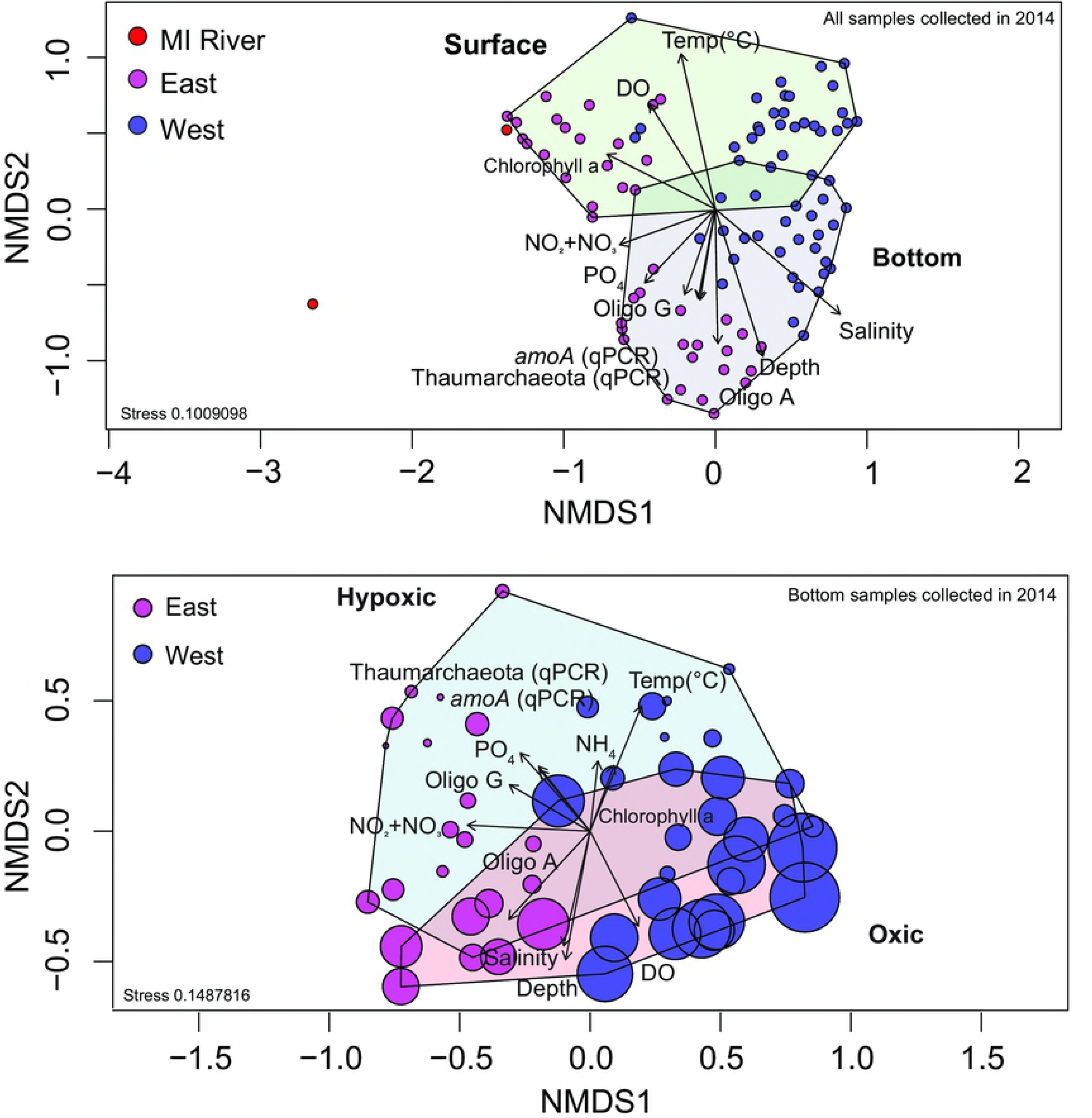
NMDS ordination of normalized 16S rRNA gene iTag sequence data. (A) NMDS ordination of normalized 16S rRNA gene iTag sequence data for all Y14 surface and bottom samples. Mississippi River samples (MI) are in red **(A)**, samples east of the Atchafalaya River (AR) are in magenta and samples west of the AR are in purple. **(B)** NMDS ordination of normalized 16S rRNA gene iTag sequence data of Y14 bottom samples only. Bubble sizes represent DO concentrations e.g. larger bubbles indicate higher DO concentrations. All environmental variables shown as vectors were significantly (corrected p-values ≤ 0.05) correlated with an NMDS axis for both **(A)** and **(B)**.

To analyze beta diversity in Y14 bottom water hypoxic verses oxic conditions, surface samples were excluded and bottom water samples were examined using NMDS ordination (Fig 2B). An adonis test revealed distinct microbial communities in hypoxic samples as compared to oxic water samples (Fig 2B) (adonis R^2^= 0.09 and p-value ≤ 0.05, beta-dispersion F= 0.60 p-value ≥ 0.05). Bottom samples showed significant spatial clustering east and west of the AR (adonis R^2^= 0.35 and p-value ≤ 0.05, beta-dispersion F= 0.01 p-value ≥ 0.05) (Fig 2B).

### Taxonomic and functional gene abundances in the 2014 hypoxic zone

Bacterial and thaumarchaeal 16S rRNA and archaeal *amoA* gene copy numbers (qPCR) were determined in surface and bottom water samples. Bacterial 16S rRNA gene copy numbers L^-1^ were similar in the surface (avg. 4.10 × 10^7^) and in the bottom water samples (avg. 3.73 × 10^7^). In contrast, thaumarchaeal 16S rRNA gene copy numbers L^-1^ were significantly higher in bottom water samples (avg. 2.18 × 10^7^) compared to surface samples (avg. 2.19 × 10^6^) as were *amoA* copy numbers L^-1^ (bottom avg. 3.12 × 10^7^ and surface avg. 2.96 × 10^6^) (Wilcoxon, S2 Table). When comparing hypoxic and oxic samples in bottom only samples, thaumarchaeal 16S rRNA gene copy numbers L^-1^ were significantly higher in hypoxic water samples (avg. 3.28 × 10^7^) compared to oxic water samples (avg. 8.87 × 10^6^) as were *amoA* copy numbers L^-1^ (avg. hypoxic 4.65 × 10^7^ and avg. oxic 1.32 × 10^7^) (S2 Table In surface and bottom water samples, and in bottom water only samples, thaumarchaeal 16S rRNA and *amoA* gene copy numbers L^-1^ were significantly positively correlated with each other, NO_2_+NO_3_ and PO_4_ and significantly inversely correlated with DO (S4-7 Tables). The ratio of Thaumarchaeota 16S rRNA:*amoA* gene copy number L^-1^ was one (avg.).

### Differences in the extent of hypoxia and microbial community structure between years 2013 and 2014

The total area of low oxygen in Y14 was 13,080 km^2^, compared to 15,120 km^2^ in Y13 (gulfhypoxia.net). Comparing the same hypoxic sites we sampled in Y13 and Y14 revealed that DO in Y13 samples was lower (avg. 0.62mg/L) than Y14 (avg. 1.1mg/L) and the average depth of the hypoxic zone was deeper in Y13 (17.7m) compared to Y14 (14.6m) (S4 Fig).

Ammonium, salinity and temperature were significantly different between the two hypoxic zones (Wilcoxon, S2 Table) with average NH_4_ concentrations being higher in Y13. In the Y13 hypoxic zone, the average ammonium and salinity concentrations were higher, while in the Y14 hypoxic zone (which was located at shallower depths) the average temperature was higher.

Alpha diversity, Shannon, observed (species richness) and equitability were not statistically significantly different between the years. To look at beta diversity between the years, the bottom samples collected at the same stations in Y13 and Y14 were examined using NMDS (S5 Fig), with environmental variables that were significantly correlated with NMDS axes represented as vectors (corrected p-values ≤ 0.05, S8 Table). An adonis test revealed distinct microbial communities in Y14 samples as compared to Y13 samples (S5A Fig) (adonis R^2^= 0.20 and p-value ≤ 0.05, beta-dispersion F= 1.07 p-value ≥ 0.05). Whereas Y14 showed distinct east and west clusters, Y13 did not (adonis R^2^= 0.09 and p-value ≤ 0.05, beta-dispersion F= 10.47 p-value ≤ 0.05) (S5A Fig).

Analysis of normalized iTag sequence data of bottom samples collected at the same stations in Y13 and Y14 revealed that the phyla Thaumarchaeota, Actinobacteria, Planctomycetes, Euryarchaeota and SAR406 were greater in Y13 than Y14, whereas Cyanobacteria, Proteobacteria and Bacteroidetes were greater in Y14. At the OTU level, 17 had significant normalized abundances between Y13 and Y14 (Wilcoxon). Five of the 17 OTUs were more abundant in Y14, whereas 12 OTUs were more abundant in Y13 (S6 Fig). Specifically, Thaumarchaeota, *N. maritimus* OTU4369009 (S6 Fig), a Thaumarchaeota OTU4073697 (95% similar to cultured representative *Nitrosopelagicus brevis* strain CN25) and Euryarchaeota, MGII OTU3134564 had higher normalized abundances in Y13 than in Y14. Of the five OTUs that had higher abundances in Y14, one was a Thaumarchaeota OTU1584736 (95% similar to cultured representative *Nitrosopelagicus brevis* strain CN25). The two Thaumarchaeota that were 95% similar to *Nitrosopelagicus brevis* strain CN25 had a one nucleotide (nt) base pair differentiation at 82/250 (G-A) in the 16S rRNA gene sequence.

When comparing absolute abundance data in the same hypoxic sites sampled in both years, thaumarchaeal 16S rRNA and archaeal *amoA* gene copy numbers (qPCR) per L averages were significantly higher (S4 Fig and S2 Table) in Y14 (avg. 3.31 × 10^7^ and 4.52 × 10^7^) than in Y13 (avg. 7.25 × 10^6^ and 6.87 × 10^6^). In both years, copy number per L of each gene was significantly inversely correlated with DO (Spearman’s ρ and corrected p-values in S9-12 Tables).

### Shannon entropy analysis of Nitrosopumilus 16S rRNA gene sequence data in the 2013 and 2014 hypoxic zones

Due to the high abundances of *Nitrosopumilus* in hypoxic samples in Y13 and Y14, oligotyping analysis was carried out to examine closely related *Nitrosopumilus* (all sequences annotated as such) in relationship to environmental variables. The Y13 iTag data were not analyzed in this way in our previous paper [20], so here we present CSS normalized OTU data oligotyping results for both Y13 and Y14 in the same station samples (n = 35/year). The normalized abundance of oligotype G (G in nt position 115/250) was significantly higher in abundance in hypoxic samples and more abundant in Y13 compared to Y14 (Wilcoxon, S2 Table). Oligotype G was significantly inversely correlated with DO in both Y13 and Y14 (S9-12 Tables). Conversely, oligotype A (A in nt position 115/250) was lower in hypoxic samples in both years (Fig 3). Oligotypes T or C at nt position 115/250 were lower, reaching maximal abundances of 6.45% and 0.29%, respectively.

### Thaumarchaeota AOA microbial species co-occurrence patterns in the 2013 and 2014 hypoxic zones

Similar to oligotyping analysis, species co-occurrence was not considered in our Y13 dataset. We therefore evaluated species co-occurrences by determining Spearman’s correlation coefficients for the same Y13 and Y14 bottom water sites (n = 35/year). For this analysis, we included only the 17 OTUs discussed above that were significantly different between the years (Wilcoxon), which included the three Thaumarchaeota OTUs; 4369009, 1584736, 4073697 (S6 Fig). Of the 17 OTUs that were significantly different between Y13 and Y14 same station samples, ten OTUs were significantly correlated at least once with one of the three Thaumarchaeota OTUs in Y13 and/or Y14 hypoxic or oxic samples (Fig 4).

**Figure 4.**
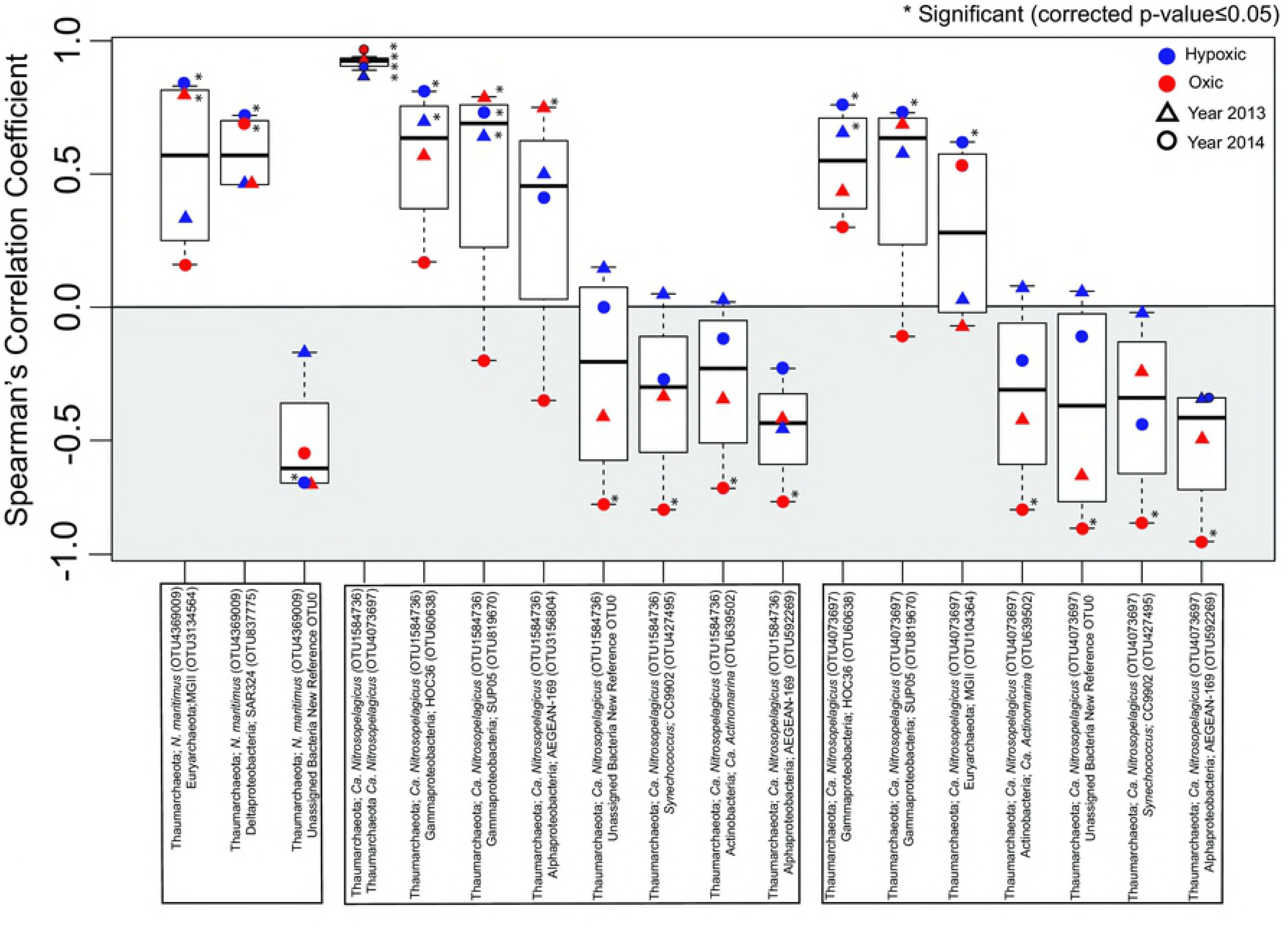
Plots representing co-occurrence (Spearman’s ρ) data for OTUs of interest. Co-occurrence (Spearman’s ρ) of Thaumarchaeota (*N. maritimus* OTU4369009, *Ca. Nitrosopelagicus* OTU1584736 and *Ca. Nitrosopelagicus* OTU4073697) and ten OTUs whose - normalized abundances were significantly different between years (Wilcoxon test with B-H corrected p-values) in the same Y13 and Y14 bottom water samples. Correlations that were significant (corrected p-values ≤ 0.05) are indicated by an *.

## Discussion

The Y14 hypoxic zone was slightly smaller in size (13,080 km^2^), closer to shore and discontinuous as compared to the Y13 hypoxic zone (15,120 km^2^) (S4 Fig). In July 2014, wind speeds reached between 10 to 20 knots blowing from the west, with higher than average wave height (1.4 meters) reported (http://www.wavcis.lsu.edu/). Further, in Y14 there was an unseasonably late (late July), above average Mississippi River discharge (avg. is × 4 × 10^5^ ft^2^/sec compared to 5 × 10^5^ ft^2^/sec in Y14), which resulted in nitrogen (NO_2_ + NO_3_) concentrations reaching a near-record high (200 μmol L^-1^) and an anomalously high phytoplankton biomass (e.g., 118 ug/L Chl *a* at the end of July) (LUMCON 2014, http://bit.ly/2wHLSk1, accessed 02/08/17) late in the hypoxic season (S1 Table and S1 Fig). These factors could have resulted in the slight DO replenishment seen between 92° W and 92.38° W, specifically along transect G, west of the Atchafalaya River (AR) (S1 Table and S1 Fig). Previous reports of wind out of the west and southwest during the summer months have correlated with a smaller hypoxic zone, which moves nutrient enhanced water masses to the east and to deeper waters, disrupting density stratification [69,70]. Therefore, a plausible hypothesis is that the variability in wind forcing resulted in the movement of water masses to the east [71]. This wind forcing could have influenced the hypoxic area to the west of the Atchafalaya River, resulting in the variability in the size and shape of the hypoxic zone between Y13 and Y14, while also potentially shaping the microbial community as seen in the east west latitude clustering in Y14, but not in Y13 (S4 and 5 Figs).

Specifically, our data revealed a significant inverse correlation between AOA and DO in the nGOM. However, in Y14 when the hypoxic zone was less extensive, and focused at the MI River mouth the normalized abundances of AOA were lower than in Y13. In Y14, although AOA were enriched in the hypoxic bottom water, after correcting for multiple comparisons, their normalized abundances (iTag) were not significantly higher in hypoxic verses oxic samples. Yet the absolute abundances of thaumarchaeal 16S rRNA and *amoA* genes were significantly higher in Y14 than in Y13 (S4 Fig). This discrepancy between iTag and qPCR suggested that the iTag primers did not discern the full breadth of Thaumarchaeota diversity in comparison to qPCR primers. To reconcile the disparity between iTag and qPCR, samples that had high Thaumarchaeota 16S rRNA and *amoA* gene copy number were further analyzed. Both MI River samples had low normalized abundances of Thaumarchaeota (less than 1%), but up to 10^5^ copies of Thaumarchaeota 16S rRNA genes/L and 10^6^ copies of *amoA* genes/L. These results could be indicative of Thaumarchaeota that are introduced to the nGOM during freshwater input, compared to Thaumarchaeota that are adapted to saline conditions introduced to the hypoxic zone via nGOM seawater in the bottom layer. Therefore, the location and physical parameters of the hypoxic zone could influence Thaumarchaeota abundance and subsequently overall microbial diversity.

Further comparison of Y13 and Y14 revealed that the *Nitrosopumilus* oligotype G was both annually abundant in the nGOM and was significantly inversely correlated with DO (Fig 3). This oligotype was differentiated by 1 bp in all 16S rRNA gene sequences annotated as *Nitrosopumilus*. Whether this polymorphism is ubiquitous in low DO adapted Thaumarchaeota is not yet known. The data did suggest that its abundance was influenced by the severity of bottom water hypoxia (expansive and deeper verses shallow and smaller), where abundances were greater in Y13 (Fig 3). Thus, this *Nitrosopumilus* oligotype may be adapted to low oxygen, coastal conditions. Oligotyping has revealed ecologically important sub-OTUs in human and environmental microbiomes across environmental gradients [66,72-75]. For example, Sintes and colleagues [74] identified two groups of *amoA* oligotypes that clustered according to high or low latitude and subclustered by ocean depth, however little other oligotypic analysis has been carried out on AOA, specifically *Nitrosopumilus*. Thus, oligotyping can reveal subtle nucleotide variations within AOA. Whether a low DO adapted AOA oligotype that was dominant in coastal hypoxic samples are ecologically consequential remains to be determined.

Microbial species co-occurrences relationships can reveal community patterns, and facilitate hypothesis generation regarding abiotic influences on random and non-random patterns [61-64]. In our study, co-occurrence analysis of OTUs that were significantly different between bottom water sites in Y13 and Y14 revealed that increasing normalized abundances of *N. maritimus* OTU4369009, particularly in Y13, potentially perturbed co-occurrence relationships with other OTUs, specifically MGII Euryarchaeota OTU3134564 and Deltaproteobacteria OTU837775 (putatively identified as a SAR324 by SILVA) (Fig 4). Previous studies reported that MGII have an aerobic photoheterotrophic lifestyle [76-78] with higher abundances in the euphotic zone compared to depths below the euphotic zone [28,79-81]. Whereas Thaumarchaeota, closely related to the *N. maritimus*, increase in abundance with depth [17,18,28,79,82]. Therefore, if high thaumarchaeal abundances in the hypoxic zone are not met with higher MGII abundances, potential metabolic linkages would be decoupled, which could be ecologically significant. It has been reported that some SAR324 have the ability to oxidize hydrogen sulfide [83] which could, in theory, mean that when hypoxic conditions prevail and AOA continues to draw down DO, SAR324 could oxidize sulfide that may flux from the sediments resulting in detoxification of bottom water. During hypoxic conditions when *N. maritimus* OTU4369009 abundances are high, co-occurrence with SAR324 abundances are weakened (Fig 4) and sulfide oxidation by SAR324 may not keep pace, resulting in a deteriorating environment beyond low DO.

The two Thaumarchaeota (OTU1584736 and OTU4073697) that were significantly different between the years were significantly positively correlated with each other in both hypoxic and oxic samples in both years (Fig 4). These two OTUs are 95% similar to the *N. Brevis* CN25 [84], an AOA that has been shown to produce N_2_O enrichment cultures [85], similar to *N. maritimus* [86,87]. AOA have been reported to be the primary source of N_2_O in the surface ocean [85] and it has been shown that decreasing oxygen concentration could influence the production of N_2_O by AOA [85,86,88–90]. In the shallow nGOM water column, Walker JT and colleagues [91] reported that nitrification was the primary source of N_2_O during peak hurricane season, consistent with the results of Pakulski (2000) [25]. Therefore, the three Thaumarchaeota OTUs that we show to be abundant in the n GOM hypoxic zone of both years may contribute to N_2_O production, a potent greenhouse gas [92,93], which can flux from ocean to the atmosphere when the water column is mixed by tropical storm activity [91].

Additionally, a decoupling between the two-step transformation of ammonium to nitrate (which are carried out by distinct groups of microorganisms) has previously been reported (e.g. [94,95], including in the nGOM hypoxic zone [19]. In our dataset nitrate oxidizing bacteria (NOB) were undetectable, thus the annual increase in AOA in the nGOM, and the lack of co-occurrence with NOB suggested that nitrite may accumulate, as shown by Bristow and colleagues [19] in this expansive hypoxic zone. This metabolic decoupling, leading to NO_2_ accumulation during the summer months when the nGOM hypoxic zone develops, is consequential given the toxicity of nitrite [96].

## Conclusion

Collectively, this dataset supports that the nGOM hypoxic zone can serve as a hotspot for AOA and the that the normalized abundance of Thaumarchaeota 16S rRNA (iTag) and the absolute abundance (qPCR) of Thaumarchaeota 16S rRNA and archaeal *amoA* gene copy numbers can reflect the extent of bottom water hypoxia. There are few reports describing abundant Thaumarchaeota in shallow coastal environments; most of which represent polar environments [97-101]. Our findings of an increase in Thaumarchaeota in the hypoxic nGOM are consistent with several studies that have reported an increase in abundance of Thaumarchaeota in low oxygen marine environments [18–20,33–36], however this is first known dataset that has sampled the water column microbial community of the shelfwide nGOM hypoxic zone in two consecutive years. Future studies that determine the ecological implications of an AOA hotspot, their co-occurrences and the potential impact on biogeochemical cycling, such as the contribution to N_2_O production and their role in ocean deoxygenation, which is intensifying in a changing climate [1,2,102–104] are needed.

## Author Contributions

LGC, NNR and JCT collected samples. LGC carried out DNA extractions, library preparation and sequencing and statistical analysis of the data. OUM designed the project and did the bioinformatics. LGC and OUM wrote the manuscript.

## Acknowledgements

We thank the science and vessel crews of the R/V *Pelican* and Louisiana Universities Marine Consortium for their valuable shipboard and onshore support.

## Supporting Information

**S1 Fig. (A)** Sample map of the 52 stations sampled during the Y14 nGOM shelfwide cruise, in which bottom water status is indicated (red circles indicates oxic stations while blue circles indicates hypoxic stations). (**B)** DO concentrations from the bottom samples collected.

**S2 Fig. (A)** NMDS ordination of normalized 16S rRNA gene iTag sequence data for all samples collected in year 2014 grouped by surface and bottom. The seventeen bubble plots represent the same NMDS plot with normalized abundances of the OTUs that were statistically significantly different (Wilcoxon) between surface and bottom samples depicted by bubble size, where larger bubble size represents higher normalized abundances. The symbol * represents OTUs that were statistically significantly inversely correlated with dissolved oxygen (Spearman correlation; B-H corrected p-value ≤ 0.05).

**S3 Fig. (A)** NMDS ordination of normalized 16S rRNA gene iTag sequence data for the all bottom samples collected in year 2014 where bubble size represents DO concentration. The other sixteen NMDS bubble plots represent the normalized abundances of the OTUs that were statistically significantly different (Wilcoxon) between hypoxic and oxic samples. Larger bubble size represents higher normalized abundances. The symbol * represents OTUs that were statistically significantly inversely correlated with dissolved oxygen while the other OTUs are significantly positively correlated with DO (Spearman correlation; B-H corrected p-value ≤ 0.05).

**S4 Fig.** Plots of DO concentrations, relative abundances of Nitrosopumilus OTU4369009 16S rRNA genes (iTag), Thaumarchaeota 16S rRNA and amoA gene copy number/L (qPCR) for bottom water samples collected at the same sites in Y13 and Y14.

**S5 Fig. (A)** NMDS ordination of normalized 16S rRNA gene iTag sequence data for the same samples collected in Y13 and Y14 with environmental variables, qPCR data and oligotype data shown as vectors. All environmental variables represented as vectors on the NMDS were significantly correlated (corrected p-value ≤ 0.05) with an NMDS axis. **(B)** The same NMDS ordination depicting oxygen concentration (DO) as bubble size e.g. larger bubbles indicate higher DO concentrations.

**S6 Fig. (A)** NMDS ordination of normalized 16S rRNA gene iTag sequence data for the same samples collected in Y13 and Y14 where bubble size depicts oxygen concentration. The other seventeen NMDS bubble plots represent the normalized abundances of the OTUs that were statistically significantly different (Wilcoxon) between the years where larger bubble size represents higher normalized abundances. The symbol * represents OTUs that were statistically significantly inversely correlated with DO in Y14, and the symbol # represents a significant inverse correlation with DO in Y13 (Spearman correlation; B-H corrected p-value ≤ 0.05).

**S1 Table.** 2014 nGOM hypoxic zone sample metadata and in situ chemistry.

**S2 Table.** Wilcoxon B-H corrected p-values for diversity statistics and environmental variables for all datasets.

**S3 Table.** Operational taxonomic unit (OTU) table for all samples collected in 2014.

**S4 Table.** P-values (B-H corrected) for Spearman’s correlation coefficients for Y14 surface and bottom samples.

**S5 Table.** Spearman’s Correlation coefficients (ρ) for Y14 surface and bottom samples.

**S6 Table.** P-values (B-H corrected) for Spearman’s correlation coefficients for Y14 bottom samples only.

**S7 Table.** Spearman’s Correlation coefficients (ρ) for Y14 bottom samples only.

**S8 Table.** Envfit B-H corrected p-values for NMDS ordinations.

**S9 Table.** P-values (B-H corrected) for Spearman’s correlation coefficients for Y13 same samples only.

**S10 Table.** Spearman’s Correlation coefficients (ρ) for Y13 same samples only.

**S11 Table.** P-values (B-H corrected) for Spearman’s correlation coefficients for Y14 same samples only.

**S12 Table.** Spearman’s Correlation coefficients (ρ) for Y14 same samples only.

